# Ataxin-2 Disordered Region Promotes Huntingtin Protein Aggregation And Neurodegeneration In Drosophila Models Of Huntington’s Disease

**DOI:** 10.1101/2021.10.01.462783

**Authors:** Joern Huelsmeier, Emily Walker, Baskar Bakthavachalu, Mani Ramaswami

## Abstract

The Ataxin-2 (Atx2) protein contributes to the progression of neurodegenerative phenotypes in animal models of amyotrophic lateral sclerosis (ALS), type 2 spinocerebellar ataxia (SCA-2), Parkinson’s Disease (PD) and Huntington’s Disease (HD). However, because the Atx2 protein contains multiple separable activities, deeper understanding requires experiments to address the exact mechanisms by which Atx2 modulates neurodegeneration (ND) progression. Recent work on two ALS models, C9ORF72 and FUS, in *Drosophila* has shown that a C-terminal intrinsically disordered region (cIDR) of Atx2 protein, required for assembly of ribonucleoprotein (RNP) granules, is essential for the progression of neurodegenerative phenotypes as well as for accumulation of protein inclusions associated with these ALS models. Here we show that the Atx2-cIDR also similarly contributes to the progression of degenerative phenotypes and accumulation of Huntingtin protein aggregates in *Drosophila* models of HD. Because Huntingtin is not an established component of RNP granules, these observations support a recently hypothesised, unexpected protein-handling function for RNP granules, which could contribute to the progression of Huntington’s disease and, potentially, other proteinopathies.

## INTRODUCTION

Neurodegenerative diseases (ND) differ in clinical pathologies. For instance, ALS primarily causes the degeneration of motor neurons, spinocerebellar ataxias primarily affect cerebellar Purkinje cells and Huntington’s disease affects cortical and striatal neurons (Vonsattel *et al*. 1985; Seidel *et al*. 2012; Braak *et al*. 2013). However, the presence of pathological protein aggregates, nuclear, cytoplasmic or even extracellular, is a shared feature of most neurodegenerative diseases (Ross and Poirier 2004; Ramaswami *et al*. 2013; Soto and Pritzkow 2018). Most notably: extracellular Amyloid-ß (A-ß) plaques as well as neurofibrillary tangles of hyper-phosphorylated Tau protein occur in Alzheimer’s disease patients (Duyckaerts *et al*. 2009; Peng *et al*. 2020); prion protein filaments are found in prion diseases (Prusiner 1991); and α-synuclein and ubiquitin containing inclusions called Lewy bodies (LB) occur in Parkinson’s disease (Polymeropoulos *et al*. 1997; Spillantini *et al*. 1997; Luk *et al*. 2012; Henderson *et al*. 2019). Similarly, the Spinocerebellar ataxias (SCAs) caused by expansions of intrinsic poly-glutamine tracts in the causative proteins, are associated with the presence of respective mutant-protein aggregates, particularly in neurons that lie within vulnerable brain regions (Lastres-Becker *et al*. 2008; Seidel *et al*. 2012; Rüb *et al*. 2013). The intracellular location of such inclusions can vary: for instance, in SCA-3, neuronal inclusions of Ataxin-3 are mostly nuclear, while in SCA-2 inclusions of Ataxin-2 are mainly cytoplasmic (Huynh *et al*. 2000; Seidel *et al*. 2017).

A causal role of these usually amyloid-rich intracellular aggregates in disease is argued by several lines of evidence. For Alzheimer’s disease, striking genetic analyses have shown that human mutations in APP that promote the production of A-ß also promote Alzheimer’s disease and mutations that specifically reduce A-ß production confer resistance (Jonsson *et al*. 2012). A common pathway by which such aggregates potentially cause disease, is suggested by increased levels of integrated stress response (ISR) signalling in animal models of neurodegenerative disease (Moreno *et al*. 2012). Elevated levels of misfolded proteins results in activation of stress kinases, which by phosphorylating eIF2a reduce translation efficiency (Harding *et al*. 2003; Wek *et al*. 2006; Pavitt 2018). Consistent with a causal role in disease progression, restoring protein translation by blocking eIF2a phosphorylation or its consequences can delay, block or even reverse disease progression in animal models (Moreno *et al*. 2012; Sidrauski *et al*. 2015; Halliday *et al*. 2015, 2017). Therefore, although definitive clinical support for the “amyloid hypothesis,” that amyloid aggregates play major causal roles in disease, remains elusive, there is considerable interest in therapeutic strategies based on either inhibiting ISR signalling or reducing the formation of disease specific protein inclusions (Duennwald and Lindquist 2008; Shacham *et al*. 2019; Ganz *et al*. 2020).

Three main lines of evidence implicate ribonucleoprotein (RNP) granules in the formation of disease-associated protein aggregates in ALS and Frontotemporal Dementia (FTD). First, several RNP granule components including TDP-43 are enriched in ALS/FTD associated neuronal inclusions (Hart *et al*. 2012; Kim *et al*. 2013; Gopal *et al*. 2017; Markmiller *et al*. 2018). Second, mutations in different RNP granule components have been found in ALS/FTD patients (Elden *et al*. 2010; Ramaswami *et al*. 2013; Taylor *et al*. 2016). Third, many pathogenic mutations occur in assembly-prone intrinsically disordered regions (IDRs) commonly found in RNA-binding proteins and act to promote aggregate formation (Guo *et al*. 2011; Kim *et al*. 2013).

Several analyses have shown that the RNP-granule protein Ataxin-2 has multiple functions, being required for miRNA activity, translational repression, translational activation, mRNA polyadenylation and RNP-granule assembly (Nonhoff *et al*. 2007; Mccann *et al*. 2011; Lim and Allada 2013; Zhang *et al*. 2013; Sudhakaran *et al*. 2014; Bakthavachalu *et al*. 2018; Inagaki *et al*. 2020). Ataxin-2 protein is also required for pathological aggregate formation and neurodegeneration in animal models of ALS/FTD (Elden *et al*. 2010). Thus, knockdown of Ataxin-2 via antisense oligonucleotides delays the maturation and recruitment of TDP-43 to RNP granules (stress granules) *in vitro*, reduces pathologies and extends lifespans in a TDP-43 mouse model of ALS (Becker *et al*. 2017; Scoles *et al*. 2017). Studies in *Drosophila* have clarified that of Ataxin-2’s multiple activities, its specific role in RNP-granule assembly contributes to the progression of neurodegenerative phenotypes in fly models of ALS/FTD (Elden *et al*. 2010; Bakthavachalu *et al*. 2018). Thus, genetic deletion of a C-terminal intrinsically disordered region (cIDR) required for efficient formation of RNP granules results in perfectly viable adult flies, which, remarkably, show resistance to cytotoxicity in two *Drosophila* ALS models, of C9ORF72 and FUS, respectively (Bakthavachalu *et al*. 2018). Taken together with additional observations that the Atx2-cIDR is also required for the aggregation of FUS protein and C9-associated dipeptide repeats in cell culture, these results support a model in which RNP granules, perhaps by concentrating aggregation-prone proteins within membrane-less organelles, enhance the efficiency with which initial pathogenic aggregates associated with ALS/FTS can be nucleated (Becker and Gitler 2018).

A more recent study of Huntington’s Disease (HD) in *Drosophila* showed a connection between polyQ expanded Huntingtin (Htt) and Ataxin-2 (Xu *et al*. 2019b). In Huntington’s disease, pathogenic expansions occur within an endogenous polyQ domain starting at amino acid 17 of Htt. Via proteolytic processing and aberrant splicing, an N-terminal region of the Htt protein is separated from the full-length protein. This fragment containing expanded poly-Q domains is found in cellular aggregates (Landles *et al*. 2010; Sathasivam *et al*. 2013; Peskett *et al*. 2018). Expression of pathogenic forms of Htt-polyQ in *Drosophila* resulted in the formation of Htt-polyQ aggregates most easily visualized in cells of the eye imaginal disc, death of adult photoreceptors or, if appropriately expressed, in death of cells that control the circadian clock in *Drosophila* (Jackson *et al*. 1998; Romero *et al*. 2008; Weiss *et al*. 2012; Schilling *et al*. 2019; Xu *et al*. 2019b). Remarkably RNAi mediated knockdown of Atx2, in clock cells expressing Htt-polyQ, reduced both Htt-polyQ aggregate formation and cytotoxicity in these cells (Xu *et al*. 2019b).

Given that Htt is not an established component of neuronal RNP granules, we were interested in examining whether Atx2’s RNP-granule forming ability was also required for its role in promoting Htt-polyQ induced protein aggregation and neurodegeneration. We report that removal of the Atx-2 cIDR not only protects against photoreceptor degeneration in two *Drosophila* HD models but also greatly reduces the Htt-polyQ aggregate size and number in eye imaginal disc cells. We document these observations, which are consistent with recent reports indicating overlapping pathways and similarities between ALS and HD cellular pathologies (Freibaum *et al*. 2015; Zhang *et al*. 2015; Grima *et al*. 2017). We discuss why these results may support a model in which RNP granules serve a secondary general function in sequestration of misfolded cellular proteins (Al-Ramahi *et al*. 2007b; Lessing and Bonini 2008; Ganassi *et al*. 2016; Mateju *et al*. 2017).

## MATERIALS AND METHODS

### Drosophila husbandry and fly stocks used

*Drosophila* stocks were maintained at 25°C in cornmeal agar. Experimental crosses were carried out at 25°C and 60% humidity with a 12h light dark cycle. The GMR-Gal4 (BDSC 1104) and UAS-htt-ex1-Q97 (BDSC 68417) lines were obtained from the Bloomington Drosophila Stock centre (Indiana). UAS-mRFP-mHtt-Q138 and UAS-mRFP-mHtt-15Q were a gift from Troy Littleton (Weiss *et al*. 2012). UAS-TDP-43-wt and UAS-TDP-43-M337V were a gift from Paul Taylor (Ritson *et al*. 2010). *atx2-ΔcIDR-GFP* was previously published (Bakthavachalu *et al*. 2018).

### Immunofluorescence and confocal imaging

Wandering 3^rd^ instar larvae were collected from respective genetic crosses and eye imaginal discs were processed for immunostaining as previously described (Sudhakaran *et al*. 2014). Each immunostaining experiment was reproduced in at least three independent biological replicates: for each replicate, discs dissected from multiple larvae were individually examined.

Primary antibodies used were: rabbit anti-RFP (Invitrogen R103367,) at 1:1000, rabbit-anti-Rasputin (Aguilera-Gomez *et al*. 2017) 1:1000, rabbit-anti-Caprin (Papoulas *et al*. 2010) at 1:1000, rat-anti-Elav-7E8A10 (Developmental Studies Hybridoma Bank) 1:300 and chicken-anti-Ataxin-2 (Bakthavachalu *et al*. 2018) at 1:500. Secondary antibodies were used at 1:1000 dilution: Alexa Fluor®488 goat anti-Chicken (Invitrogen A11039), Alexa Fluor®488 goat anti-Rabbit (Invitrogen A11078), Alexa Fluor®488 goat anti-Rat (Invitrogen A11006), Fluor®555 goat anti-Rabbit (Invitrogen A21428).

Prepared eye-imaginal discs were mounted in Vectashield Mounting Medium (Vector Labs, H-1000) on microscope slides and imaged on a Zeiss LSM880 confocal microscope.

### Eye degeneration

To assess eye degeneration *GMR-Gal4* was to respective UAS-effector lines to appropriately model degenerative disease. To examine the role of the Atx2-cIDR in promoting degeneration in these contexts, double balanced *GMR-Gal4*; *atx2-ΔcIDR-GFP* flies were crossed to UAS-effector lines that either included *atx2-ΔcIDR-GFP*, if the effector transgene (e.g UAS-Htt-138Q-RFP) was on the second chromosome, or recombined with *atx2-ΔcIDR-GFP* if the effector transgene (e.g UAS-mRFP-mHtt-Q138, UAS-mRFP-mHtt-15Q, UAS-htt-ex1-Q97) was on the third chromosome.

Fly eyes were scored within 24 hours of eclosion and every 10 days thereafter until day 50. Eye phenotypes were scored into 3 categories. Normal eyes, that did not have any discoloration or visible roughness. Mild phenotypes, represent eyes that have local or wide spread discoloration, roughness or glossiness without necrotic areas. Strong phenotypes represent eyes that additionally exhibit necrotic areas identified by black spots.

The number of flies scored per genotype are listed in Figure 1 (A-F) and Supplementary Figure 2 (A’’-E’’). Bar graphs in the panels of Figure 1 (A’’’-F’’’) and Supplementary Figure 2 (A’’-E’’) represent the percentage of flies corresponding to each phenotype and dead flies at the respective day. Both male and female flies were examined; for logistical reasons, scoring was not performed blind to genotype.

**Figure 1:**
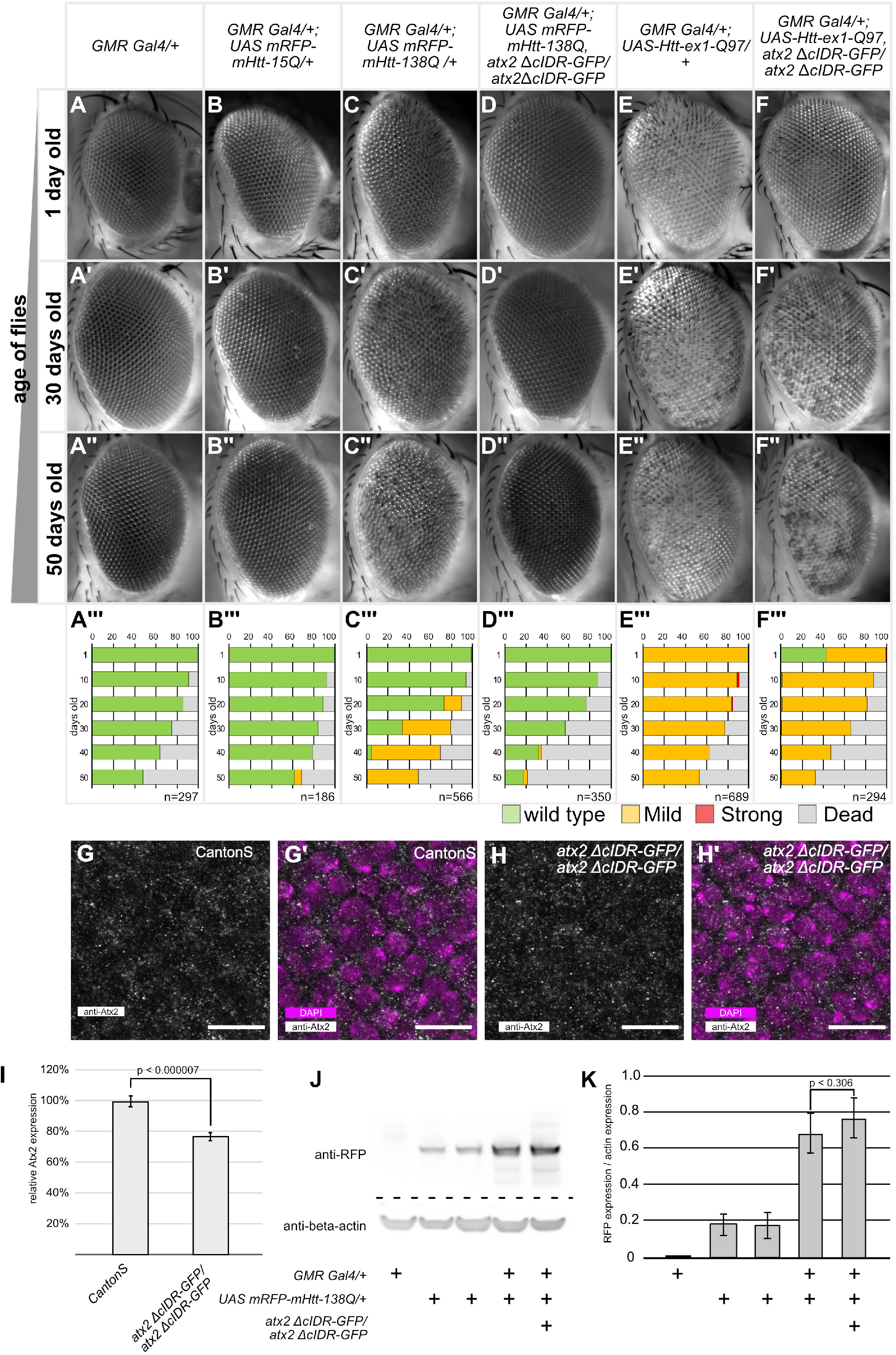
Atx2-cIDR-GFP is required for Htt-polyQ induced neurodegeneration. A-F’’: Representative images of *Drosophila* eyes taken at the indicated time points (1 day old, 30 days and 50 days old, female flies). Genotypes are indicated at the top of each column. Inserts are quantifications of phenotypes for the corresponding genotype and age (male and female flies). A’’’-F’’’ quantification of phenotypes observed at 10-day intervals between 1-day and 50-days after eclosion. Green bars indicate percentages of wild-type eyes, yellow bars a mild phenotype with patches of discoloration or rough surface visible, red bars are eyes with necrotic spots and grey bars percentage of dead flies. Number (n) for each genotype at day 1 is indicated in the insert. G-H’) Confocal images of eye imaginal discs of 3^rd^ instar *Drosophila* larvae, posterior is up. G/H) Tissue stained with anti-Atx2 antibody (grey). G’/H’) Tissue stained with anti-Atx2 antibody (grey) and DAPI (magenta). G/G’) Wild type Canton S, H/H’) homozygous *atx2 ΔcIDR-GFP*. Sub cellular localisation of wild type Atx2 is similar to Atx2 ΔcIDR-GFP. Scale bars 10μm. I) Quantification of fluorescence intensity of Atx2 antibody reactivity, normalized to fluorescence intensity in CantonS. Three biological replicates per genotype, at least 5 discs were analysed per replicate. Expression levels of *atx2 ΔcIDR-GFP* are significantly lower than wild-type Atx2 (2-tailed Student’s t-Test, p=0.0000069). Error bars SEM. J) Representative Western blot of head lysates from 1 day old flies, 10 heads per lane. Genotypes are indicated below image. Immuno-reactivity to anti-RFP (top) and anti-beta-actin (bottom). K) Quantification of the expression levels of UAS mRFP-mHtt-138Q as measured by Western-Blot immune-reactivity to anti-RFP. mRFP-mHTT expression levels are normalised to the corresponding beta-actin loading control. Three independent replicates were carried out. Genotypes are indicated below the bar graph. Expression levels of mRFP-mHtt-138Q in *GMR-Gal4/+;UAS-mRFP-mHtt-138Q, atx2-ΔcIDR-GFP/ atx2-ΔcIDR-GFP* are not significantly different from the levels in *GMR-Gal4/+;UAS-mRFP-mHtt-138Q* flies (one-tailed Student t-test, p=0.305). Error bars are SEM.

### Eye image acquisition

In addition to flies used to score phenotypes, a small batch of flies was aged separately. Flies of this cohort were sacrificed and used for image-acquisition. At specific ages flies were transferred into a 1.5ml micro-centrifuge tube and frozen at 20°C. For imaging, flies were pinned with insect needles to a Sylgard plate. In order to image the whole compound eye, flies were oriented with one eye clearly visible. Automated Z-stack images were taken on a Zeiss AxioImager Z1 with a 10x lens, with a stack distance of 18μm. To generate the final images, image-stacks were “focus stacked” using ImageJ (Schneider *et al*. 2012), followed by orientation and cropping of the images. In all images, anterior is to the right, dorsal to the top.

### Protein Extraction from Drosophila Heads

For each genotype 40 fly heads of 1 day old flies were collected on ice, and transferred into a microcentrifuge tube containing pre-cooled 40μl extraction buffer (20 mM HEPES, pH 7.5, 100 mM KCl, 5% glycerol, 10 mM EDTA, 0.1% Triton, 1 mM dithiothreitol, 0.5 mM phenylmethylsulfonyl fluoride, 20 mg/mL aprotinin, 5 mg/mL leupeptin, 5 mg/ mL pepstatin A). Heads were ground on ice with a pestle for 30 seconds followed by brief centrifugation at 4°C. This was repeated three times. After the last repeat samples were centrifuged for 5 minutes at 4°C, the supernatant was collected into a new microcentrifuge tube. The volume of each tube was adjusted to 40μl with Extraction buffer. 4μl 10x Sample Reduction Agent (Invitrogen, B0009) and 10μl 4x LDS Sample Buffer (Invitrogen B0007) were added to the samples and heated for to 70°C for 10 minutes. Samples were then stored at -80°C until use.

*Drosophila* head lysates, corresponding to 10 heads per sample, were separated by SDS gel electrophoresis and subsequently transfered using the iBlot 2 System (Invitrogen, IB21001S) according to the manufacturer’s instructions. Western blots were performed using rabbit anti-RFP (abcam ab62341, 1:5,000) and mouse anti-beta-actin (Proteintech 6008-1, 1:5000). Secondary antibodies conjugated with HRP from Jackson Immunoresearch were used at 1:10,000 dilution.

### Image analysis

Western blot images were analysed using the gel densitometry tool in ImageJ (Schneider *et al*. 2012). Expression of mRFP-mHtt-Q138 was calculated in relation to the expression of the beta-actin loading control. The experiment was independently repeated in triplicate.

For immunofluorescence experiments, mHtt-polyQ granules were identified using the Analyse Particle function in ImageJ. In short, z-projections of confocal images of 3^rd^ instar larvae eye discs were cropped to include the entire GMR-Gal4 expressing domain as defined by the anti-elav staining. Images were blurred (Gaussian blur, Sigma =2), thresholded and despeckled to create a binary image for particle detection. The Analyse Particle function of imageJ was used to identify aggregates (parameters: size=50-Infinity pixel circularity=0.70-1.00). The number of discs per analysed genotype is listed in Fig. 2, the average size of the area analysed per disc was 15,665 μm^2^.

**Figure 2:**
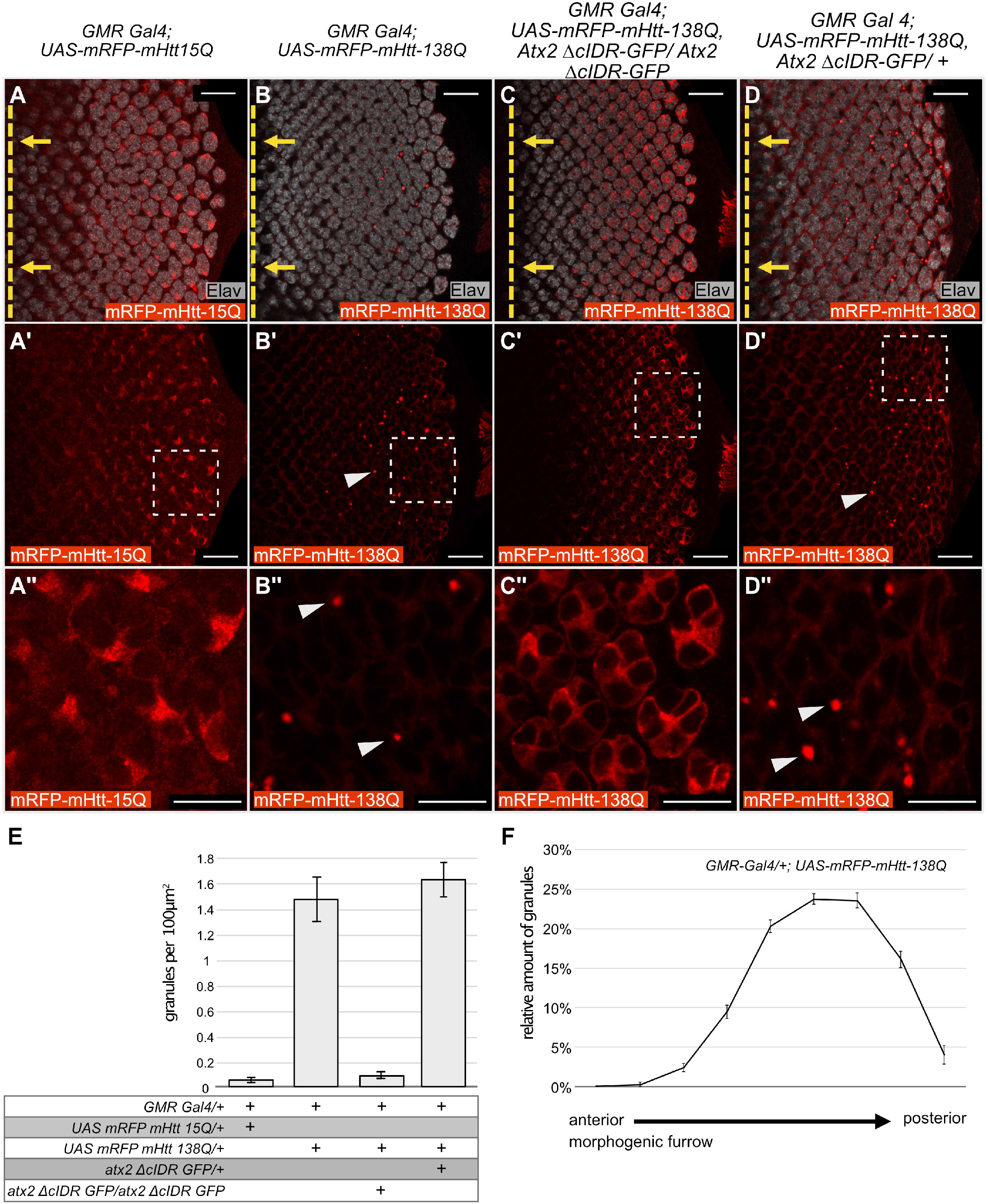
Atx2-cIDR is required for aggregation of Htt-polyQ in *Drosophila* eye discs. A-D’’) Confocal images, single optical sections, of eye imaginal discs from *Drosophila* third-instar larvae expressing different *UAS-mRFP-mHtt-polyQ* constructs with *GMR-Gal4*. Red: anti-RFP to visualize mHtt-polyQ. Grey: anti-Elav to identify neurons (photoreceptor cells) in the developing disc. Posterior is to the right, the position of the morphogenic furrow is indicated by a dashed yellow line in A-D. Yellow arrows indicate the direction of relative movement of the morphogenic furrow. A’’-D’’ are 4x magnifications of the indicated areas in A’-D’ respectively. No aggregation is observed for *UAS-mRFP-mHtt-15Q* (A-A’’). When over expressed mRFP-mHtt-138Q assembles into aggregates (arrowheads in B’,B’’). This aggregation is dependent on the Atx2 cIDR, as aggregates do not form when the cIDR is not present (C-C’’). The effect of the Atx2 cIDR is dominant as aggregates of mRFP-mHtt-138Q still form when wild type Atx2 is present (arrow heads in D’,D’’). Scale bars (A-D’) 20μm, (A’’-D’’) 10μm. E) Quantification of the amount of Huntingtin aggregates in A-D. Bar graphs represent the amounts of aggregates observed per 100μm^2^. Genotypes are indicated below the graph. From left to right: 1) *GMR Gal4/+; UAS-mRFP-mHtt-15Q /+* (n=8) 2) *GMR Gal4/+; UAS mRFP-mHtt-138Q /+* (n=8) 3) *GMR Gal4/+; UAS mRFP-mHtt-138Q, atx2 ΔcIDR-GFP/atx2 ΔcIDR-GFP* (n=9) 4) *GMR Gal4/+; UAS mRFP-mHtt-138Q /atx2 ΔcIDR-GFP/+* (n=3). Error bars are SEM. F) Distribution of mRFP-mHtt-138Q aggregates along the anterior-posterior axis of larval eye imaginal discs starting at the morphogenic furrow (left hand side of the graph) to the posterior edge of the imaginal disc (right hand side of the graph) (n=8). Error bars are SEM. The number of aggregates increases with distance to the morphogenic furrow.

Atx2 expression levels were calculated based on confocal image stacks collected from eye imaginal discs of either CantonS or *atx2-ΔcIDR-GFP* 3^rd^ instar larvae. To ensure the antibody signal between genotypes was not affected by deviations in the staining protocol between reaction vials, we carried out the staining protocol for both genotypes in the same vials. When scanning we distinguished between the two genotypes based on a dsRed expression only present in the Bolwig nerve of *atx2-ΔcIDR-GFP* flies. We measured the mean fluorescent intensity of Atx2 across multiple levels of the disc using ImageJ. Measurement was limited to the GMR-Gal4 expression domain. The domain was approximated based on the antibody staining for anti-elav. Three independent repeats of the experiment were carried out, with six imaginal discs analysed per replicate and genotype. For each imaginal disc the relative mean fluorescent intensity in relation to the mean fluorescent intensity of Atx2 signal in the Canton-S control was calculated.

## RESULTS

### Atx2-cIDR is partially required for the progression of mHtt-polyQ induced degeneration

One of the hallmarks of Huntington’s disease is the presence of intracellular aggregates of polyglutamine containing Htt protein fragments. Expression of CAG expanded Htt in the *Drosophila* eye leads to photoreceptor degeneration and disorganization of the ommatidial units of the compound eye (Romero *et al*. 2008). We first confirmed these previous observations and then, to determine if the RNP-granule forming activity of *Drosphila* Atx-2 contributes to Htt-polyQ induced degeneration, tested if degenerative phenotypes were altered if the granule-assembly domain of Atx2 was deleted.

We expressed UAS mRFP-mHtt-polyQ constructs in the eye via the *GMR-Gal4* driver, which induces expression of UAS effectors in all cells posterior to the morphogenetic furrow of the eye imaginal discs (Freeman 1996). Eyes of flies expressing *UAS-mRFP-mHtt-15Q* (a non-pathogenic 15Q repeat) with *GMR-Gal4* were indistinguishable from control flies and did not exhibit any macroscopic degradation (Figure 1 A-B’’’). In contrast, expression of the CAG expanded *UAS-mRFP-mHtt-138Q* construct led to a clearly visible and progressive degeneration of the compound eye with age (Figure 1 C, C’’’). A small subset of flies developed mild localised discoloration of the ommatidia at day 20 after hatching (Figure 1 C’’’). By day 50, this discoloration is visible in almost all experimental flies (Figure 1 C’’,C’’’). A similar phenotype can be observed in flies expressing *UAS-Htt-ex1-Q97*. Here the phenotype is already apparent from the first day of hatching (Figure 1 E-E’’’). We then asked how these phenotypes were affected by the removal of the cIDR domain of Atx2, which is required for RNP-granule assembly. Western blotting confirmed that levels of mRFP-mHtt-138Q were similar in experimental *GMR-Gal4/+;UAS-mRFP-mHtt-138Q, atx2-ΔcIDR-GFP/ atx2-ΔcIDR-GFP* and *GMR-Gal4/+;UAS-mRFP-mHtt-138Q* in *Drosophila* head extracts (Figure 1, J/K, one-tailed Student t-test, p=0.306). The specificity of the immunoreactivity was confirmed as no RFP protein is detected in *GMR-Gal4* driver controls. Surprisingly, we detected expression of the *UAS-mRFP-mHtt-138Q* allele in absence of the *GMR-Gal4* driver. Therefore we anticipated that any phenotypic difference would be due to the change in activity of the Atx2 protein.

To test whether mHtt-polyQ induced degeneration of the eye is dependent on the Atx2-cIDR, we crossed either *UAS-Htt-ex1-Q97* or *UAS-mRFP-mHtt-138Q* into genetic backgrounds where the Atx2-cIDR was deleted via CRISPR mediated genome engineering (Bakthavachalu *et al*. 2018). In both cases we observed a suppression of the degenerative phenotypes (Figure 1: D-D’’, F-F’’). In the case of mRFP-mHtt-138Q we could not observe any discoloration in fly eyes even 50 days after eclosion (Figure 1, D-D’’). The difference in phenotype is significant at day 30 and 50 (chi-squared test p < 0.00001) but not at day 1 (chi-squared test p=0.265). The phenotype induced by *UAS-Htt-ex1-Q97* in freshly eclosed flies is suppressed by the removal of Atx2-cIDR (Figure 1, E-E’’, F-F’’) and is significantly different from the phenotype observed in one day old *GMR;UAS-Htt-ex1-Q97* flies (chi-squared test p<0.00001). However, for *UAS-Htt-ex1-Q97* this suppression is no longer evident after 10 days (chi-squared test at 30 days p=0.1134, at 50 days p=0.1241) (Figure 1, F’-F’’).

To address the possibility that suppression of mHtt-polyQ degeneration is due to reduced expression levels of the *atx2-ΔcIDR-GFP* mutant, we compared the levels of Atx2 in eye imaginal discs of homozygous *atx2-ΔcIDR-GFP* and CantonS flies. Homozygous *atx2-ΔcIDR-GFP* flies do not contain any wild-type Atx2, hence immunoreactivity to the Atx2 antibody reports presence of the Atx2-ΔcIDR-GFP protein only (Bakthavachalu *et al*. 2018). We found that in both genotypes Atx2 is similarly distributed in the cells of the eye imaginal disc (Fig. 1 G-H’). We noted that the level of Atx2 quantified using immunofluorescence in homozygous *atx2-ΔcIDR-GFP* flies is slightly (∼20%) but significantly lower when compared to the expression level in Canton-S flies (Figure 1, I, 2-tailed Student’s t-Test, p=0.0000069). However, this small reduction, which may indicate a subtle increase in diffuse, less easily measured cytoplasmic labelling, seems unlikely to account for the observed phenotypic differences. As previously shown, the immunoreactivity we observe is specific to the Atx2 protein (Singh *et al*. 2021). Taken together, these observations indicate that as for *Drosophila* ALS models, progressive cytotoxicity in *Drosophila* Huntington’s models is facilitated by the Atx2-cIDR.

An unexpected caveat to this arises from our observation is that the survival rates of flies expressing mHtt-polyQ in *atx2-ΔcIDR-GFP* background are reduced when compared to wild-type controls (Figure 1, inserts in A-F’’, Suppl. Fig. 1). Thus, we cannot exclude the possibility that the above experiments inadvertently compare eyes of a healthy subset of experimental flies, for instance with low levels of Htt-polyQ, with a broader range of control flies. While the caveat needs to be noted, we note that prior work on Atx2 has shown that the protein itself is required for Htt-polyQ induced toxicity in clock neurons (Xu *et al*. 2019b). Furthermore, suppression of the phenotype induced by *UAS-Htt-ex1-Q97* in the *atx2-ΔcIDR-GFP* background is observed well before excessive lethality becomes an issue. Thus, our results are indicative of a likely role for the cIDR promoting Htt-polyQ induced toxicity.

To further examine this working hypothesis, we tested whether the formation or stability of Htt-polyQ aggregates observed in Htt-polyQ expressing eye imaginal disc cells were affected by deletion of the cIDR.

### Atx2-cIDR is required for the aggregation of mHtt-polyQ protein

To test whether the Atx2-cIDR is required for aggregation of mHtt-polyQ proteins *in vivo*, we stained and visualized mRFP-mHtt-138Q inclusions in *Drosophila* eye imaginal discs. In control flies expressing the short glutamine-repeat fragment *UAS-mRFP-mHtt-15Q* with *GMR-Gal4*, mHtt-15Q protein appeared diffuse and cytoplasmic in photoreceptor progenitor cells and no inclusions could be seen (Figure 2, A-A’’/ E, 0.04 granules per 100 μm^2^). In contrast, in eye-imaginal disc cells expressing the pathogenic polyQ expansion fragment *UAS-mRFP-mHtt-138Q*, we observed obvious cytoplasmic inclusions of the mutant protein (Figure 2, B-B’’/ E, 1.01 granules per 100 μm^2^). The presence of mRFP-mHtt-138Q aggregates was predominantly found in the posterior half of the imaginal disc (Figure 2 F). This argues for increased aggregation over time, as cells near the optic stalk (posterior) are older as compared to cells close to the morphogenetic furrow (anterior) (Ready *et al*. 1976). This increased aggregation over time is broadly consistent with the late onset of the macroscopically visible phenotype observed in adult *Drosophila* eyes (Figure 1 C-C’’). When expressed in larvae homozygous for the Atx2-cIDR deletion, the mRFP-mHtt-138Q protein distribution is strikingly changed to a diffuse and cytoplasmic pattern, without any apparent aggregates or inclusions (Figure 2, C-C’’/ E A, 0.07 granules per 100 μm^2^). Additional control experiments show mHtt-138Q-RFP inclusions in control flies heterozygous for *atx2-ΔcIDR-GFP* (Figure 2 D-D’’/ E, 1.29 granules per 100 μm^2^) that still carry one wild-type copy of Atx2. The lack of mHtt-138Q-RFP aggregates in the absence of the Atx2-cIDR argues that Atx2-dependent granules contribute to mRFP-mHtt-138Q aggregate accumulation and indicate a shared mechanism for aggregate formation across at least two different classes of neurodegenerative diseases. It is interesting that while Htt aggregates can form as early as the third-instar larval stage, degeneration of the adult eye is only obvious around 30 days after eclosion for mRFP-mHtt-138Q and 10 days after eclosion for Htt-ex1-Q97. Additional experiments are required to understand how Htt aggregates are connected to cell death. One possibility is that aggregates are tolerated in developing imaginal discs, and cause cell death only when their levels or their negative effects on cell metabolism build up over days. Alternatively, it may be that eye defects we score under a dissection microscope do not reveal low levels of cell death that may occur, but be invisible to external inspection, in younger flies.

### mHtt-polyQ aggregates do not sequester stress granule markers

The ALS associated C9-dipeptide repeat and FUS proteins are known to localize substantially to RNA stress granules in cultured cells and/or to inclusions that contain stress granule markers like TDP-43 or Ataxin-2 *in vivo* (Liu-Yesucevitz *et al*.; Bentmann *et al*. 2012; Bakthavachalu *et al*. 2018; Chew *et al*. 2019) To test whether Htt-polyQ granules also similarly sequester stress-granule markers, we examined Htt-polyQ aggregates *in vivo* and in eye imaginal discs for colocalization with stress granule markers. None of the stress-granule marker proteins tested, including Atx2, Caprin and Rasputin (G3BP), showed colocalization with mHtt-138QRFP in imaginal disc cells (Figure 3,A-C). These experiments do not provide any additional insight into how Atx2 facilitates mHtt-polyQ aggregate formation but also do not, on their own, rule out a transient role for stress granules in early stages of mHtt-138QRFP aggregate formation.

**Figure 3:**
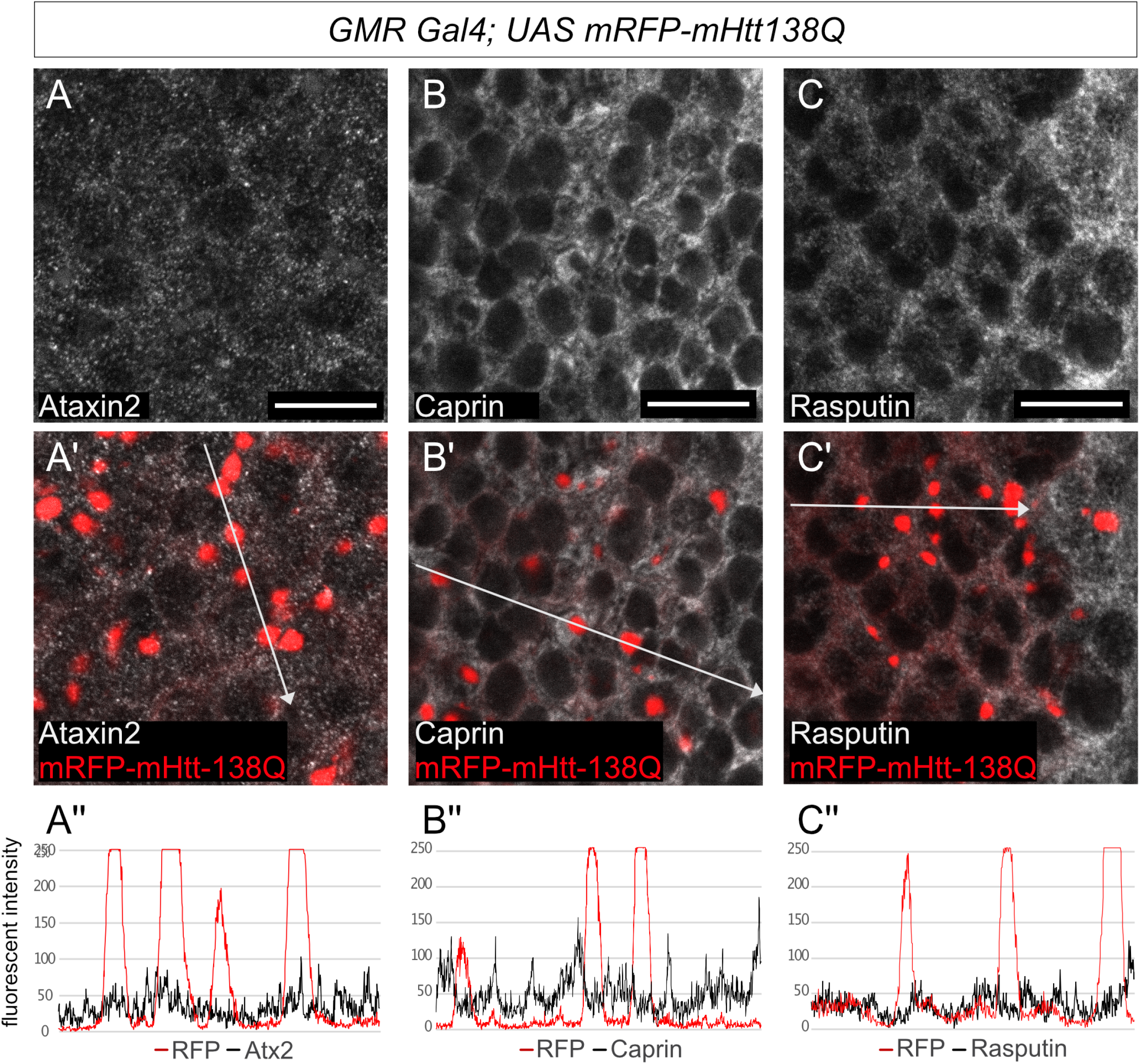
Ectopic polyQ Huntingtin aggregates do not colocalise with stress granule markers. A-C’ Confocal images of *Drosophila* eye discs from third-instar larvae expressing *UAS-mRFP-mHtt-138Q* with *GMR Gal4*. red: mRFP-mHtt-138Q shown by RFP-fluorescence, grey: A-A’ anti-Atx2, B-B’ anti-Rasputin, in C-C’ anti-Caprin. Posterior is to the right. Scale bars 10μm. A’’-C’’ Fluorescence intensity plots of the antibody signals measured along the white arrow in A’-C’ respectively. None of Atx2, Caprin nor Rasputin co-localise with mRFP-mHtt-138Q aggregates.

## DISCUSSION

We previously showed that the cIDR of Atx2 is required for the progression of degenerative phenotypes in two ALS models in the fly: C9orf72 di-peptide and FUS (Bakthavachalu *et al*. 2018). In the course of this study, we performed additional experiments to allow us to extend this conclusion to *Drosophila* TDP-43 models for ALS. Overexpression of each UAS-TDP-43-wt and UAS-TDP-43-M337V in the fly eye leads to degeneration of the ommatidia with different degrees of severity (Ritson *et al*. 2010). The TDP-43-M337V mutation, which results in the substitution of residues in a phylogenetically conserved region of TDP-43, is linked to familial cases of ALS (Sreedharan *et al*. 2008). While previous experiments have demonstrated that TDP-43 induced cytotoxicity is dependent on Atx2 (Ritson *et al*. 2010) our current analysis additionally shows that degeneration of the fly eye in TDP-43 models is facilitated by the Atx2-cIDR (Supplementary Figure 2 and legend).

Given that the proteins encoded by several ALS genes associate with stress granules, a role for RNP granules in promoting the formation of ALS-associated inclusions containing RNP granule components can be logically rationalized (Bakthavachalu *et al*. 2018; Becker and Gitler 2018). But could Ataxin-2 and its role in RNP-granule assembly also be relevant to other forms of neurodegeneration? Previous research in *Drosophila* has demonstrated the role of Ataxin-2 in the progression of multiple poly-glutamine-repeat diseases. Modulation of Ataxin-2 levels affects the severity of neurodegenerative phenotypes in *Drosophila* models of SCA1 and SCA3. In both disease models reduction of Ataxin-2 levels act neuroprotective whereas increased Ataxin-2 levels enhance the degenerative phenotypes (Al-Ramahi *et al*. 2007a; Lessing and Bonini 2008). Ectopic expression of human Htt-polyQ protein has previously been shown to induce granules in the cytoplasm of larval photoreceptor cells. Interestingly in adult flies Htt-polyQ protein predominantly re-locates to the nucleus of photoreceptor neurons with increased age (Jackson *et al*. 1998). A recent study on Huntington’s Disease showed that overexpression of different mutant polyQ Huntington transgenes in circadian clock neurons induces circadian arrhythmicity and aggregation of Htt-polyQ in small ventral lateral neurons (sLNVs) as well as reduction in cell numbers indicative of cytotoxicity (Xu *et al*. 2019a, 2019b). Reduction of Atx2 via RNAi knockdown alleviates both of these effects (Xu *et al*. 2019b). As removal of the Atx2-cIDR has a similar effect in larval photoreceptor cells as the knock down of Atx2 in adult clock neurons, it is likely, that the formation of further aggregates and their subsequent re-localisation to the nucleus in adult flies is also inhibited. However the precise activities of Atx2 required to promote degeneration or inclusion formation in clock neurons remains unclear.

Here we report the novel observation that the Atx2-cIDR is required for progression of neurodegenerative phenotypes as well as protein inclusion formation in fly Huntington’s disease models. This observation, which suggests a role for Atx2-dependent macromolecular assembles in HD progression, is difficult to explain because not only is Htt not obviously a component of known Atx2 containing RNP granules, but Htt-polyQ inclusions are also not known to contain additional RNP granule components. Thus, these data point to less obvious cell biological mechanisms that connect Atx2 mediated RNP granules to Htt-polyQ proteinopathy. We suggest three broad mechanisms to explain these observations.

### Model 1. Huntingtin assembles transiently into an undefined Atx2-dependent macromolecular complex

It is possible, that the granule-forming properties of Atx2-cIDR are not limited to stress granules, but can be extended to other phase separating complexes as well. There are reports that upon heat stress, endogenous Htt protein rapidly forms granules, termed Huntington Stress Bodies (HSB). These granules are reversible and may not colocalise with classical processing body and stress granule markers after heat shock (Nath *et al*. 2015). Thus, there is a possibility that a new class of RNP granule or non-RNP complex exists in which Htt can be concentrated, whose assembly may be dependent on Atx2. Loss of the Atx2-cIDR may prevent assembly of this complex, and by eliminating a structure in which Htt-polyQ is concentrated, thereby reduce opportunities for the protein to aggregate. In this context, an interesting recent study in *Drosophila* concludes that both the expanded poly-Q domain and flanking sequences impact on cellular distribution and pathology of mutant Htt, suggesting that a specific type of aggregate, and not-aggregation alone, drive neurodegeneration in HD (Chongtham *et al*. 2020).

### Model 2. Misfolded Huntingtin is dynamically concentrated in stress granules

It is possible that misfolded proteins with no function in translational control are also transiently concentrated and can assemble within stress granules. Consistent with this, recent studies in mammalian cells have shown that stress granules may serve an additional function in protein handling (Ganassi *et al*. 2016; Mateju *et al*. 2017). Time-lapse microscopy has demonstrated that a pathogenic, misfolded form of superoxide dismutase (SOD) can associate with a subset of stress granules, which first become less dynamic and then gradually evolve into pure SOD inclusions (Mateju *et al*. 2017). Defective Ribosomal Products (DRiPs) also accumulate in stress granules and promote an aberrant state, which is resistant to RNAses (Ganassi *et al*. 2016). These observations can be rationalized as follows. Granules being held together by disordered domain interactions, as facilitated by Atx2, may provide an environment in which proteins that are misfolded under conditions of stress can be stabilized, perhaps refolded and returned to the cytoplasm. Thus, SGs have a role in normal physiological protein handling (Alberti *et al*. 2017; Alberti and Hyman 2021). However, pathogenic and assembly prone forms of misfolded proteins when concentrated in SGs may efficiently form amyloid-rich aggregates that exclude native SG proteins. In this scenario, Atx2 cIDR deletion would inhibit SG assembly and reduce inclusion body formation by eliminating the microenvironment in which such pathogenic proteins are transiently concentrated. Although we do not observe the proposed transient colocalization of Htt-polyQ with SG markers in our experiments, there are previous observations consistent with such a model. In particular, after ER stress, Htt-polyQ has been reported to co-localise with Caprin1 and G3BP1 in cultured cells (Ratovitski *et al*. 2012). In a cultured Drosophila S2 cells dsRNA screen, knock down of Rasputin, the *Drosophila* homolog of G3BP1/2, has been shown to reduce Htt-polyQ aggregation (Zhang *et al*. 2010).

### Model 3. Absence of the Atx2 cIDR increases the pool of available chaperones or neuroprotective proteins

Physiological stress granules contain multiple chaperones, at least some of which have been shown to regulate their assembly and disassembly (Ganassi *et al*. 2016; Jain *et al*. 2016). A potential role for these chaperones may be to prevent stable aggregation of SG components when they are highly concentrated within RNP granules. It is therefore possible that if RNP granules are present at much lower levels in cells lacking the Atx2-cIDR, then higher levels of free soluble chaperones could be available for handling unrelated protein aggregates, such as those formed by polyQ-expanded forms of Htt.

Taken together, our experiments indicate an unexpected role for the granule forming ability of Atx2 in the progression of Huntington’s Disease-related phenotypes in a *Drosophila* model. This points to the possibility that Atx2-targeting therapeutics, which are now in development for SCA2 and ALS, may have wider use for other neurodegenerative conditions. While we propose models for how Atx2-dependent assemblies may have such roles in Huntington’s disease or other proteinopathies, additional experiments will be required to test and establish these or alternative proposals to explain the phenomena reported here.

## DATA AVILABILITY STATEMENT

Strains are available upon request. The data underlying this article are available in the article and in its online supplementary material.

## ACKNOWLEDGEMENTS

We thank members of the Baskar Bakthavatchalu lab, Amanjot Singh, Jens Hillebrand and other members of the Ramaswami and VijayRaghavan labs for several useful discussions and comments on the manuscript. Thanks to Troy Littleton and Paul Taylor for *Drosophila* lines.

